# Plasmacytoid dendritic cells in the intestine preferentially produce interferon lambda at homeostasis contributing to tonic localized innate immune responses

**DOI:** 10.1101/2025.08.25.672167

**Authors:** David A. Constant, Jacob A. Van Winkle, Philip A. Norwood, Gargi Mishra, Kimberly A. Meyer, Margaret E. Laney, Shelby R. Madden, Alec Griffith, Patrick Fernandes Rodrigues, Ram Savan, Timothy J. Nice

## Abstract

The healthy intestine maintains homeostasis in part via immune responses to microbiota, which includes basal production of interferon cytokines. Previous work showed that Type III Interferon (IFN-λ) stimulates localized pockets of interferon-stimulated genes (ISGs) in the adult mouse intestinal epithelium at homeostasis that provide preemptive protection from viral pathogens. Here, we demonstrate that a major source of homeostatic IFN-λ production in the intestine is a population of epithelium-associated plasmacytoid dendritic cells (pDC). Depletion of bacterial microbiota in the intestine also reduces pDC abundance, and pDC depletion or bone marrow reconstitution with IFN-λ-deficient pDC results in reduced expression of homeostatic ISGs in the intestinal epithelium. Notably, intestinal pDC preferentially produce IFN-λ over Type I IFNs whereas splenic pDC produce more Type I IFNs. Comparison of intestinal and splenic pDC reveal tissue-specific changes in gene expression and genomic accessibility, including evidence of responses to transforming growth factor beta (TGF-β) in the intestine. Isolated gut pDC produce more IFN-λ than splenic pDC upon stimulation, and pre-treatment of a human pDC cell line with TGF-β results in enhanced production of IFN-λ upon stimulation. This study demonstrates that pDC are an important source of homeostatic IFN-λ in the intestine and defines the role of barrier cytokine TGF-β in regulating IFN types produced by pDC upon stimulation. Reprogramming of recruited pDC by tissue cytokines may have important implications for balancing effective antimicrobial responses with damaging inflammation at barrier tissues.

**One sentence summary:** This study demonstrates that pDC are an important source of homeostatic IFN-λ in the intestine and defines the role of barrier cytokine TGF-β in regulating IFN types produced by pDC upon stimulation.

## Introduction

Plasmacytoid dendritic cells (pDC) are terminally differentiated hematopoietic cells with key roles in many immune processes (Swiecki and Colonna, 2015; Leylek and Idoyaga, 2019; Ngo et al., 2024). These cells are the main source of Type I Interferon (IFN) in the initial response to viral infection or toll-like receptor (TLR) 9 ligands (Nakano et al., 2001; Björck, 2001; Asselin-Paturel et al., 2001). There have also been several reports of pDC, as well as other DC cell types, secreting Type III IFN in response to viral infection and TLR ligands (Yin et al., 2012; Lauterbach et al., 2010). A seemingly functionally distinct subset of pDC have been shown to migrate to the intestine dependent on the C-C chemokine receptors CCR2 (Swiecki et al., 2017) and CCR9 (Wendland et al., 2007), where they mediate homeostatic and inflammatory immune responses. In addition to IFN production, pDC also affect other immune processes. For example, pDC are necessary for B cell activation and subsequent antibody production during murine rotavirus (mRV) infection (Deal et al., 2013). However, it was unknown whether gut pDC played a role in regulating homeostatic IFN responses to commensal microbiota.

Production of type I and III IFNs is stimulated by direct sensing of microbial patterns through receptors such as TLRs. The specific IFN genes produced differ, dependent on the signaling pathway or cell type that is triggered (Odendall et al., 2014; Pervolaraki et al., 2017). IFN genes are differentiated into subtypes based on their binding specificity for distinct cognate cell surface receptors (Wright and Nice, 2024), which activate signaling cascades that result in production of hundreds of interferon-stimulated genes (ISGs). Type III IFN (hereafter IFN-λ) is particularly important for stimulation of ISGs within the intestinal epithelium because mature intestinal epithelial cells (IECs) express high levels of the IFN-λ receptor and are minimally responsive to type I IFN (Pott et al., 2011; Baldridge et al., 2017; Van Winkle et al., 2020; Mahlakoiv et al., 2015; Nice et al., 2015). We have previously demonstrated that there is a highly localized ISG signature in the intestine associated with the presence of intestinal microbiota and dependent on IFN-λ signaling in IECs (Van Winkle et al., 2022; Constant et al., 2025). We showed that production of IFN-λ in this context is predominantly from an immune cell source, but the specific cell type(s) responsible remained unclear.

In this study, we identify pDC as the hematopoietic cellular source of IFN-λ in the intestinal epithelium at homeostasis. These cells preferentially produce IFN-λ, dependent on detection of microbial components and recruitment to the intestine, with minimal Type I IFN production. Furthermore, we find that pDC within the intestinal epithelium are transcriptionally and epigenetically distinct from splenic pDC, including evidence of reprogramming by transforming growth factor beta (TGF-β) in the intestine. We also find that pre-treatment of a human pDC cell line with TGF-β greatly amplifies IFN-λ production. Together, these data suggest a model in which the cytokine milieu of the intestine conditions pDC to preferentially produce IFN-λ, thereby altering IFN production in a manner that is concordant with the preferential IFN-λ responsiveness of IECs at this barrier tissue.

## Results

### Plasmacytoid dendritic cells express IFN-λ but minimal type I IFN at homeostasis

Localized IFN-λ receptor activation in IECs at homeostasis elicits pockets of ISG expression, providing preemptive protection from enteric viruses. Expression of interferon-induced protein with tetratricopeptide repeats 1 (*Ifit1*) provides a reliable proxy for the ISG response as a whole (Van Winkle et al., 2022). To definitively establish the requirement of IFN-λ cytokines for this homeostatic ISG expression in IECs, we compared *Ifit1* expression in the intestines of littermate adult mice with heterozygous or homozygous deletion of both functional murine IFN-λ genes, *Ifnl2* and *Ifnl3* (Peterson et al., 2019). *Ifit1* was robustly expressed in approximately 1-3% of the villi by RNA *in situ* hybridization in heterozygous *Ifnl2/3*^+/-^ mice, consistent with prior levels in wild-type (WT) mice, but there was virtually no detected signal in *Ifnl2/3*^-/-^ mice (**Fig. 1A-B**). These data show that the IFN-λ cytokine genes are necessary for homeostatic ISG expression in the intestinal epithelium.

**Figure 1.**
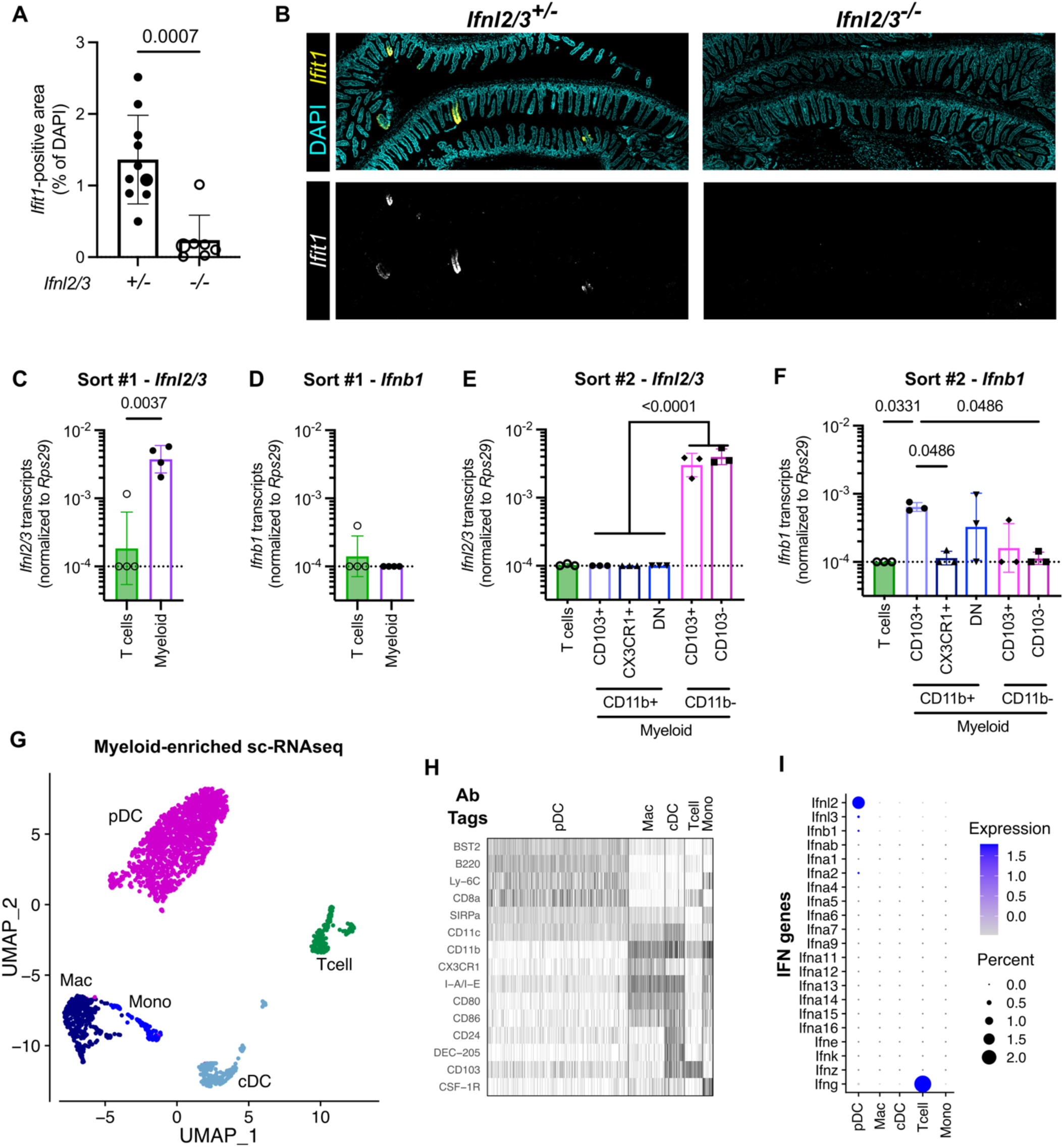
pDC express IFN-λ but minimal type I IFN at homeostasis. **A-B.** Quantitation (A) and representative images (B) showing *Ifit1* abundance in intestinal tissue from *n* = 10 *Ifnl2/3^+/-^* and *n* = 7 *Ifnl2/3^-/-^* littermate mice. Images shown in B are from mice indicated by larger data points in A. **C-F.** CD45-enriched cells from the ileal epithelium of *n* = 4 mice (C-D) or *n* = 21 mice split into pools of 7 mice each in 3 samples (E-F) were sorted for isolation of RNA and analysis of IFN gene expression by qPCR. Quantitation of *Ifnl2/3* (C, E) or *Ifnb1* (D, F) transcripts in the indicated sort populations. **G-I.** Single-cell RNAseq of cells from the ileal epithelium of *n* = 8 mice enriched for CD3^-^/MHCII^+^ cells expressing CD11c and/or CD11b. **G.** UMAP projection of cells. Clusters named based on expression of indicator genes compared to reference sets (Immgen). **H.** Heatmap of ADT counts in each cluster. **I.** Bubble plot depicting expression of all type I, II, and III IFN genes. Statistical significance by Mann-Whitney U test (A), unpaired t-test (C-D), or one-way ANOVA with Holm-Šídák’s multiple comparisons test (E-F).

We were interested in determining the specific cell lineage producing the IFN-λ responsible for stimulating homeostatic ISG expression in the intestine. Our prior work indicated that IFN-λ transcripts were detectable in CD45^+^ cells enriched from the intestinal epithelium (Van Winkle et al., 2022) and separate studies found that IFN-λ can be elicited by stimulation of dendritic cells (Lauterbach et al., 2010) or intra-epithelial T cells (Swamy et al., 2015). We first assessed homeostatic IFN-λ production by CD3^+^ intra-epithelial T cells because they are the most abundant immune cell type within the epithelial fraction. Secondarily, we sorted cells expressing CD11c or CD11b (defined here as ‘myeloid’) which should include epithelium-associated macrophages and dendritic cells (**Fig. S1A**). We isolated RNA from sorted cells for qPCR and observed significant *Ifnl2/3* transcript levels in the ‘myeloid’ subset, but undetectable transcript levels in T cells from most mice (**Fig. 1C**). In contrast to *Ifnl2/3*, the Type I IFN gene *Ifnb1* was undetectable in all subsets from most mice (**Fig. 1D**). These data indicated a myeloid cell type to be the predominant source of homeostatic IFN-λ.

To further define the cellular source of *Ifnl2/3* within the ‘myeloid’ cells above, we sub-sorted CD11b^+^ and CD11b^-^/CD11c^+^ subsets based on expression of CX3CR1 and CD103 (**Fig. S1B**), markers reported to define different mononuclear phagocytes in the small intestine (Persson et al., 2013; Farache et al., 2013). We found that nearly all *Ifnl2/3* transcripts were in the CD11b^-^/CD11c^+^ cell subset, independent of CD103 expression level (**Fig. 1E**). In contrast to *Ifnl2/3* levels, *Ifnb1* transcript levels were undetectable in CD11b^-^/CD11c^+^ subsets from most mice (**Fig. 1F**). However, we did detect *Ifnb1* transcripts in CD11b^+^/CX_3_CR1^-^ cells, which may be consistent with previous reports of homeostatic Type I IFN production by conventional DC in the gut (Stefan et al., 2020). Even so, it is notable that the magnitude of *Ifnl2/3* production was 5- to 10-fold higher than *Ifnb1* in any subset where *Ifnl2/3* was detected. This reflects the primary role of IFN-λ in stimulating homeostatic ISGs within small intestinal epithelium (Van Winkle et al., 2022).

To ensure comprehensive identification of the epithelium-associated myeloid cell subsets, we chose to pursue single-cell RNAseq as a higher-resolution method of cell type identification. CD45^+^/CD3^-^/MHCII^+^ cells from the small intestinal epithelium were sorted based on expression of CD11c or CD11b, stained with a panel of antibody-derived tags (ADTs, **Supplementary Table 1A**), and submitted for scRNAseq. A uniform manifold approximation and projection (UMAP) plot of scRNAseq data showed five distinct clusters of cells, with gene expression profiles that reflected plasmacytoid dendritic cells (pDC), classical dendritic cells (cDC), T cells (not fully excluded during sort), macrophages (Mac), and monocytes (Mono) (**Fig. 1G**). Further confidence was added to these cluster assignments based on results of the ADT labeling (**Fig. 1H**), with BST2, B220, and Ly-6C distinguishing the pDC cluster. Unexpectedly, the pDC also expressed an intermediate level of CD103, which is consistent with the staining observed in our sorting experiment above (**Fig. 1E-F, S1B**). With scRNAseq subsets defined, we next determined the expression levels of Type I, II, and III IFN genes within each subset. Relatively few cells expressed any type of IFN, but 1-2% of pDC expressed *Ifnl2* and no other cell subset expressed either *Ifnl2* or *Ifnl3* (**Fig. 1I**). There was no appreciable expression of *Ifnb1* by pDC, and none of the 18 type I IFN genes were expressed above the limit of detection within any cell subset. Type II IFN (*Ifng*) was detectably expressed by ∼2% of intra-epithelial T cells (**Fig. 1I**), which is consistent with T cells as a major source of this IFN type. Overall, these experiments show that epithelium-associated pDC are a predominant source of *Ifnl2/3*.

### pDC are recruited to the epithelium by bacterial microbiota and promote homeostatic ISG expression

We were initially surprised by the high frequency of pDC in CD3^-^ CD45-enriched intestinal epithelium (**Fig. 1G**) because pDC are not often considered in published studies of intestinal dendritic cells (e.g., Persson et al., 2013). However, two key prior studies reported pDC within the epithelial fraction at a similar frequency to our observations, and further, found that pDC recruitment to the intestine is dependent on chemokine receptors and microbiota colonization (Swiecki et al., 2017; Wendland et al., 2007). To confirm that pDC abundance is microbiota-dependent, we analyzed pDC abundance in ileal epithelium after treatment with an antibiotic cocktail (ABX) (**Fig. 2A-B**). ABX reduced pDC from an average of 1.4% to 0.4% of epithelial immune cells (**Fig. 2B**, left) whereas the proportion of CD11b^+^ myeloid cells was unchanged (**Fig. 2B**, right). As in our prior study, ABX treatment also reduced homeostatic ISG expression in the intestinal epithelium (**Fig. 2C-D**). Complementary imaging experiments to quantify pDC in the gut revealed a modest but not statistically significant decrease in pDC number after ABX treatment (**Fig. 2E**). However, when we classified pDC localization, we found that the proportion of pDC within villi was significantly lower in the ABX-treated mice (**Fig. 2F**). These data confirm prior studies of pDC recruitment to the epithelial fraction by bacterial microbiota colonization and further show that pDC recruitment correlates with homeostatic IFN-λ production.

**Figure 2.**
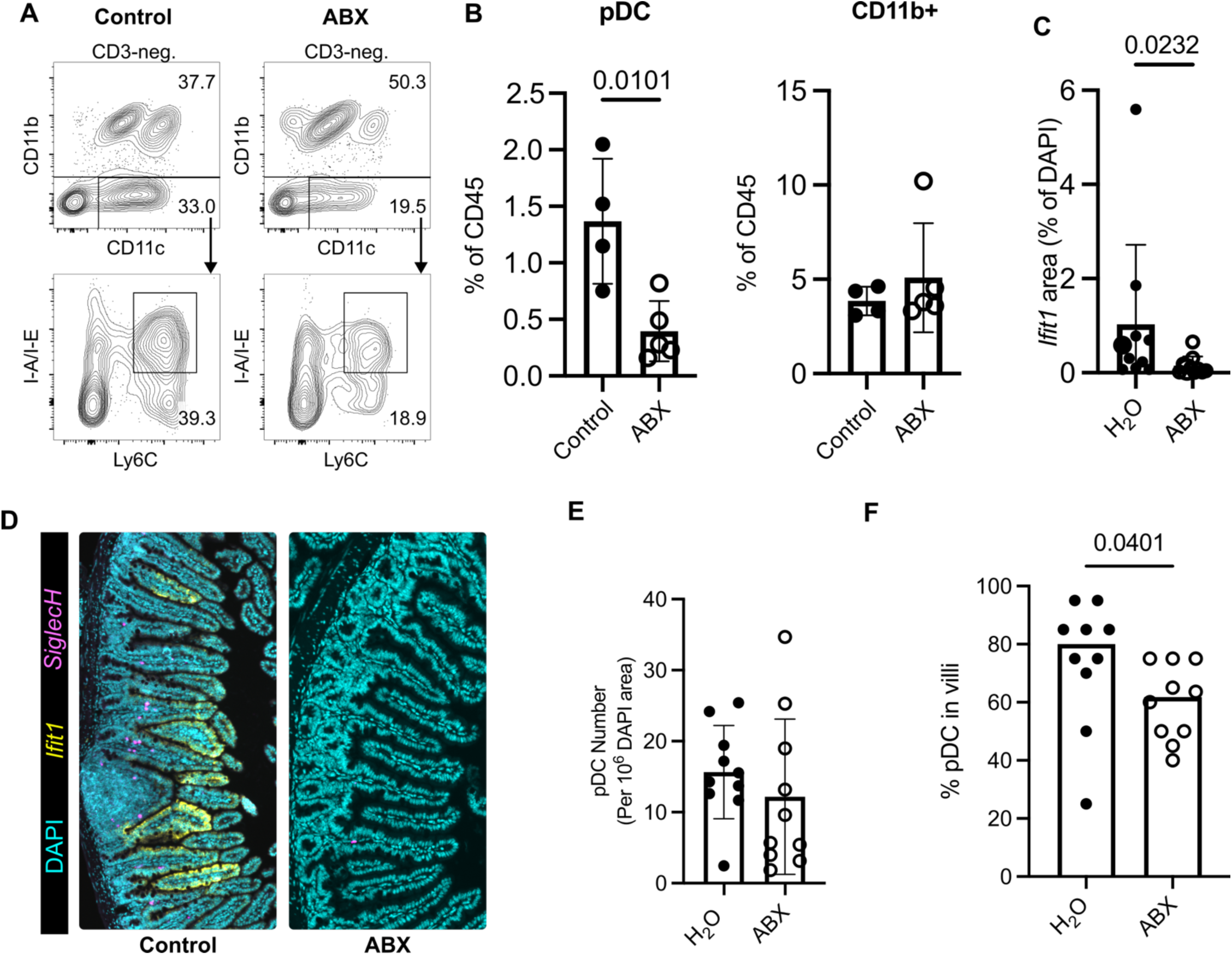
pDC are recruited to the epithelium by bacterial microbiota and promote homeostatic ISG expression. **A.** Representative flow cytometry plots, with pDC defined as CD3^-^/CD11c^+^/CD11b^-^/Ly-6C^+^/MHCII-intermediate. **B.** Proportions of pDC (left) and CD11b^+^ myeloid (right) cell subsets in mice treated with PBS (*n* = 4) or oral antibiotics (ABX, *n* = 5). **C.** Quantitation of *Ifit1* expression from RNA *in situ* hybridization performed on *n* = 10 water or *n* = 10 ABX-treated mouse intestinal Swiss rolls. **D.** Representative images showing pDC (*SiglecH*, magenta), *Ifit1* (yellow), and DAPI (cyan) from data quantified in (C). **E.** Number of pDC (as determined by *SiglecH* expression) in whole Swiss rolls. **F.** *SiglecH* puncta were used to identify pDC localized in villi versus other tissue regions from ROI manually selected at random, and the proportion of pDC in villi was analyzed. Up to 20 pDC were classified per Swiss roll, and Peyer’s patches were excluded from analysis in (F). Statistical significance by unpaired t-test (B, E) or Mann-Whitney U-test (C, F).

### pDC abundance correlates with the magnitude of homeostatic ISG responses

To directly test the requirement of pDC for homeostatic IFN-λ responses in the presence of bacterial microbiota, we depleted pDC with an anti-BST2 antibody in WT mice. This antibody has been reported to efficiently deplete mouse pDC, with minimal off-target effects (Blasius et al., 2006). As expected, this treatment reduced epithelium-associated pDC with no effect on the proportion of CD11b^+^ myeloid cells (**Fig. 3A**). BST2 antibody depletion of pDC reduced detectable *Ifit1* transcript levels in the epithelial fraction by approximately 50% relative to treatment with isotype control (**Fig. 3B**). Additional imaging analysis of pDC-depleted and control mice revealed that the *Ifit1-*positive area correlated strongly with the number of pDC detected in these tissues (**Fig. 3C-D**), linking pDC abundance with the magnitude of homeostatic IFN-λ responses.

**Figure 3.**
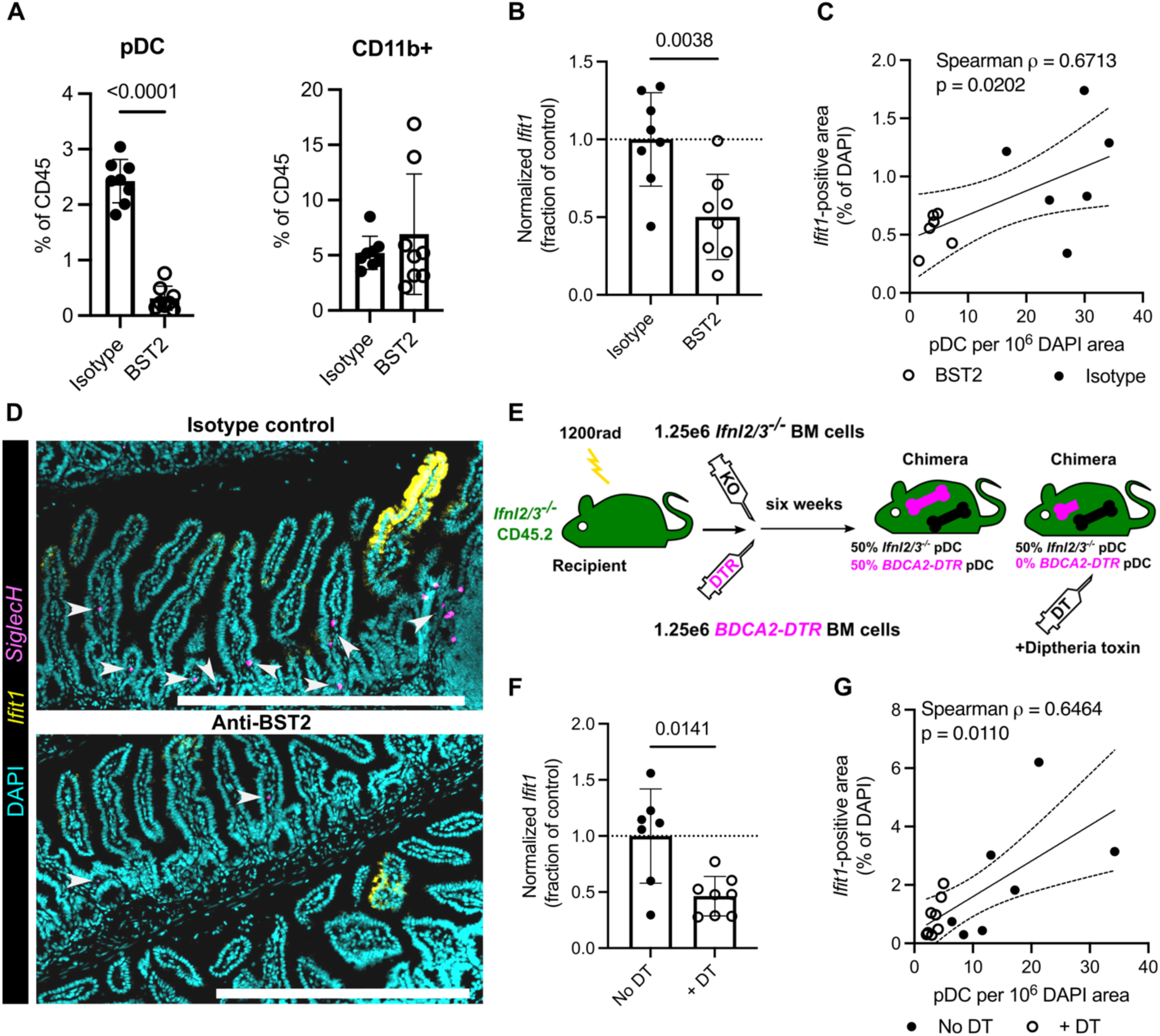
pDC abundance correlates with magnitude of homeostatic ISG responses. **A.** Proportions of pDC (left) and CD11b^+^ (right) cell subsets in epithelial fraction of mice treated isotype control or anti-BST2 pDC depleting antibody. **B.** Expression of *Ifit1* as measured by quantitative PCR from epithelial fraction of mice treated with isotype or anti-BST2 antibodies**. C.** Correlation of *Ifit1* area and pDC number from RNA *in situ* hybridization. **D.** Representative images showing pDC (*SiglecH*, magenta), *Ifit1* (yellow), and DAPI (cyan) in intestinal tissue of mice treated with isotype control and anti-BST2 antibody. *SiglecH* puncta identified as pDC are indicated by white arrowheads. **E.** Diagram of mixed bone marrow chimera generation and diptheria toxin treatment. **F.** Expression of *Ifit1* as measured by quantitative PCR from stripped epithelial fraction of mixed bone marrow chimeras with (*n* = 8) and without (*n* = 7) diptheria toxin treatment. **G.** Correlation of *Ifit1* area and pDC number from RNA *in situ* hybridization. All experiments were performed at least twice. Data in panels A-B and C-D are from separate experiments. Statistical significance by unpaired t-test (A-B), Spearman correlation (C, G), or Welch’s t-test (F).

To further test the cellular source of IFN-λ cytokines, we performed bone marrow transplant experiments using *Ifnl2/3^-/-^* mice. First, we generated reciprocal bone marrow chimeras by transferring *Ifnl2/3^-/-^* CD45.2 bone marrow or WT CD45.1 bone marrow into congenic or isogenic recipients (**Fig. S2A**). We analyzed the expression of *Ifit1* by qPCR and imaging in the small intestines of these chimeric mice six weeks later. As expected, the homeostatic ISG response was significantly higher in control WT◊WT chimeras than *Ifnl2/3^-/-^*◊*Ifnl2/3^-/-^* chimeras (**Fig. S2B-D**). Analysis of reciprocal transplants revealed that *Ifnl2/3^-/-^*◊WT chimeras had significantly greater *Ifit1* expression than *Ifnl2/3^-/-^*◊*Ifnl2/3^-/-^* chimeras, whereas WT◊*Ifnl2/3^-/-^* had an intermediate phenotype (**Fig. S2B-C**). Flow cytometric analysis of cell subsets indicated that relative proportions of T cells, pDC, and other CD45^+^ cells were roughly equivalent among all four chimeric groups (**Fig. S2E**). We also observed complete chimerism for donor cells among pDC (**Fig. S2F**), but a substantial proportion of intra-epithelial T cells were resistant to radiation (**Fig. S2G**). These data suggested that a radioresistant non-pDC cell type could be a source of homeostatic IFN-λ. However, we noted that the overall area of *Ifit1* signal was greater in WT◊WT chimeras than in WT mice not subjected to bone marrow transplant (compare **Fig. S2C** to **Figs. 1A, 2C, 3C**), which may reflect increased IFN-λ production associated with recovery from bone marrow transplant. Taken together, these data suggested that a radioresistant non-pDC cell type can amplify homeostatic IFN-λ production in the context of bone marrow transplantation.

To test the contribution of pDC-derived IFN-λ and remove the contribution of radioresistant cell types, we set up mixed bone marrow chimeras with *Ifnl2/3^-/-^* and *BDCA2-DTR* mice. *BDCA2-DTR* mice have a transgenic diptheria toxin receptor (DTR) expressed by the human *BDCA2* promoter allowing rapid, specific, and effective depletion of pDC by diptheria toxin (DT) administration (Swiecki et al., 2010). For these experiments, all recipients were *Ifnl2/3^-/-^* and were given 50% *Ifnl2/3^-/-^* and 50% *BDCA2-DTR* bone marrow (**Fig. 3E**). Resultant chimeras were then administered DT or sham six weeks later. We observed a roughly 50% reduction in homeostatic *Ifit1* expression by qPCR in the epithelial fraction of DT-treated mice (**Fig. 3F**), and imaging analyses revealed a strong correlation between pDC number and *Ifit-*positive area (**Fig. 3G**). Together with preceding figures, these data implicate microbiota-dependent recruitment of pDC to the intestine and their specific production of *Ifnl2/3* as a component of homeostatic IFN-λ signaling in the small intestine.

### IFN type preference of intestinal pDC differs from lymphoid tissue pDC and correlates with TGF-β signature genes

Plasmacytoid dendritic cells are best known for their capacity to produce Type I IFN but have been primarily studied within lymphoid tissues and may be shaped by peripheral cues (Contractor et al., 2007). Because our scRNAseq data suggest that gut pDC preferentially produce IFN-λ, we wanted to further compare pDC from gut with counterparts from lymphoid tissues. We therefore integrated our data with public scRNAseq datasets of pDC from spleen and bone marrow (Rodrigues et al., 2018; Valente et al., 2023) (**Fig. 4A-B; Supplementary Table 1B**). UMAP analysis of the integrated data revealed relatively contiguous distribution, with umap_1 largely differentiating bone marrow pDC from gut and spleen pDC, as reported in the original publication of these data by Rodrigues et al (**Fig. 4A**). Based on expression of marker genes from prior studies, we defined four sub-clusters: canonical pDC, bone marrow (BM) pDC, pDC-like cells, and ISG-expressing pDC (**Fig. 4C-D**). Notably, most cells in the gut dataset could be defined as canonical pDC with only a minority of cells clustering with related pDC-like cells (**Fig. 4C**). Most gut pDC and spleen pDC overlapped within the canonical pDC cluster (**Fig. 4A**). These data suggested that gut pDC are broadly similar to spleen pDC and less similar to BM pDC or pDC-like cells.

**Figure 4.**
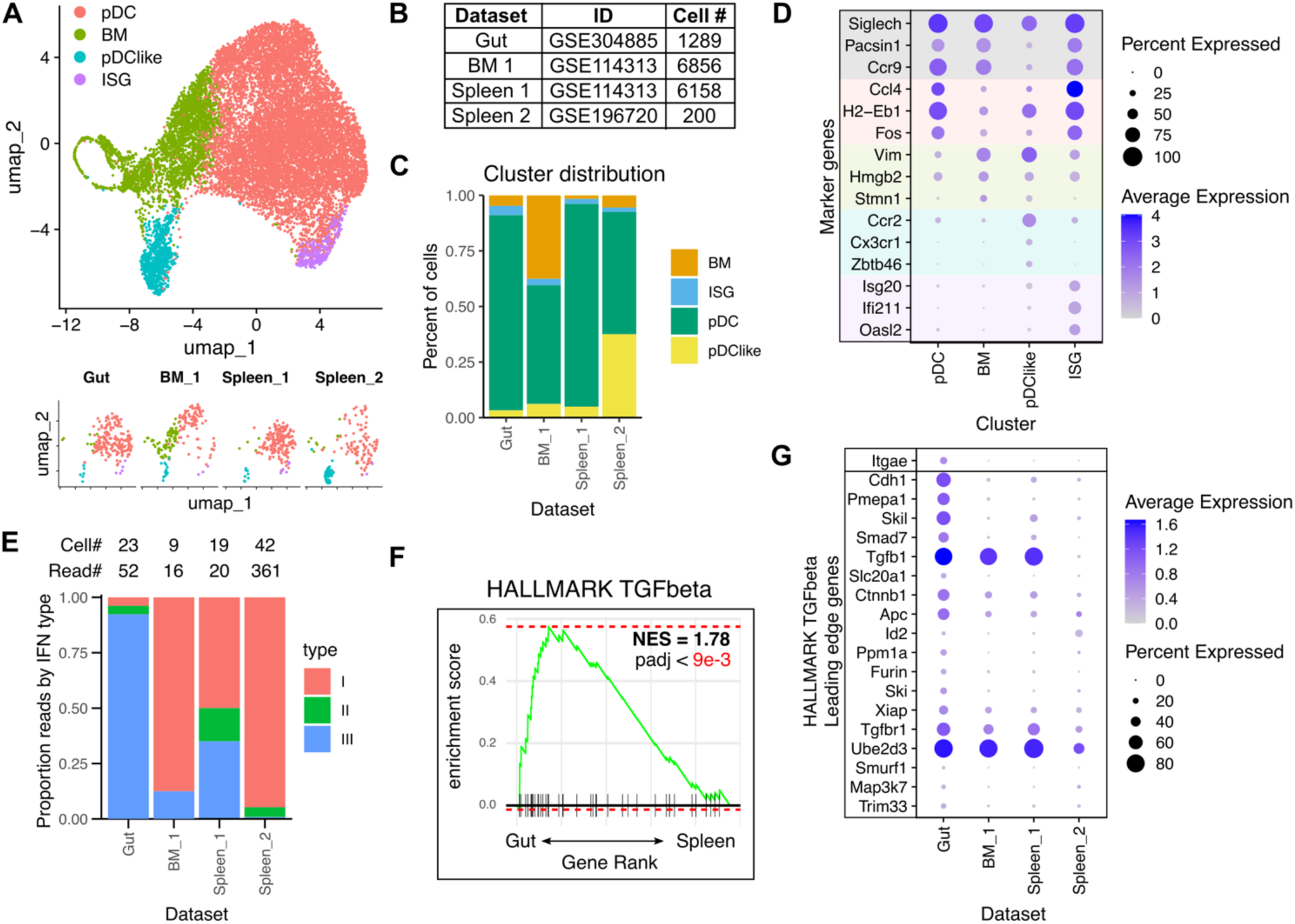
IFN type preference of intestinal pDC differs from lymphoid tissue pDC and correlates with TGF-β signature genes. Single-cell RNAseq data from this study was integrated with previously published scRNAseq datasets containing spleen and bone marrow-derived pDC. **A.** Data were integrated, and cells were clustered by UMAP. UMAP projections are shown below for each dataset individually downsampled to *n* = 200 cells to facilitate visual comparisons of cell distribution among clusters. **B.** Identifiers and cell numbers for data integrated in (A). **C.** Distribution of cells from each dataset in each cluster. **D.** Dot plot showing expression of marker genes identified by source publications for clusters, with each dataset downsampled to *n* = 200 cells. **E.** Proportion of reads belonging to type I, II, or III IFN genes in each dataset. Numbers at top indicate total number of cells expressing at least one IFN read and total number of IFN reads in each dataset. **F.** TGF-β HALLMARK pathway gene set enrichment analysis for gut vs. spleen cells in the pDC cluster. **G.** Dot plot showing expression of *Itgae* and leading-edge TGF-β HALLMARK pathway genes in each dataset.

To specifically compare IFN production among the integrated pDC datasets, we analyzed the number of cells producing detectable IFN transcripts and classified read counts by IFN type. Few cells were identified with detectable IFN reads, likely because these pDC were isolated under homeostatic conditions (**Fig. 4E**). Nonetheless, it was striking that gut pDC predominantly expressed IFN-λ (Type III IFN) while bone marrow and spleen pDC mainly expressed Type I IFNs (**Fig. 4E**).

To better understand differences between gut and spleen cells within the canonical pDC cluster, we performed HALLMARK pathway gene set enrichment analysis (**Supplementary Table 1C**). We found that TGF-β signaling was the top HALLMARK pathway enriched in gut pDC (**Fig. 4F**), with gut pDC expressing more TGF-β-stimulated genes such as *Cdh1*, *Pmepa1*, *Skil*, and *Smad7* (**Fig. 4G**). CD103 (integrin alpha E; *Itgae*) is not included within the TGF-β signaling HALLMARK pathway but is nonetheless a well-known TGF-β target gene (Bain et al., 2017) and was detected by flow cytometry and ADT labeling in our gut pDC isolation (**Fig. 1**). We found that *Itgae* was virtually undetectable in any dataset except gut pDC (**Fig. 4G**, boxed at top). Together, these data support that gut pDC are broadly similar to canonical spleen pDC but are distinguished by a TGF-β signature and preferentially produce IFN-λ at homeostasis.

### IFN type preference of intestinal pDC is associated with differential chromatin accessibility at SMAD2 binding sites

With evidence that pDC in the gut have altered IFN preference and TGF-β signature gene expression relative to splenic pDC, we were interested to investigate the underlying regulation of genomic accessibility. We therefore sorted pDC from gut or spleen (**Fig. 5A, Supplemental Fig. 3A**) and performed ATAC-seq. Motif analysis of consensus peaks in all pDC samples using the HOMER program (Heinz et al., 2010) indicated enrichment of motifs for transcription factors known to play a role in pDC development (e.g. SpiB, PU.1, E2), highlighting a shared developmental trajectory of gut and spleen pDC. Identification of differentially accessible regions between gut and spleen pDC indicated that there were 6,159 regions more accessible in gut, and 3,518 regions more accessible in spleen (**Fig. 5B**). Motif analysis within these differentially accessible regions using HOMER showed that gut-associated peaks contained a motif similar to the SMAD2 binding site, a key protein in the TGF-β signaling cascade (**Fig. 5C**). Additionally, a known SMAD2 motif was highly enriched in gut-associated peaks, and these motifs were generally close to the center of these peaks (**Fig. 5D**) suggesting active regulation by TGF-β signaling. Indeed, genes within the HALLMARK TGF-β pathway, along with *Itgae,* were overrepresented among loci with ATACseq peaks enriched in gut pDC (**Fig. 5E**), and many of these genes had increased expression in gut pDC (**Fig. 4G, 5E**, bolded genes). Together, these data suggest regulation of pDC by tissue-specific signaling factors like TGF-β in the intestine.

**Figure 5.**
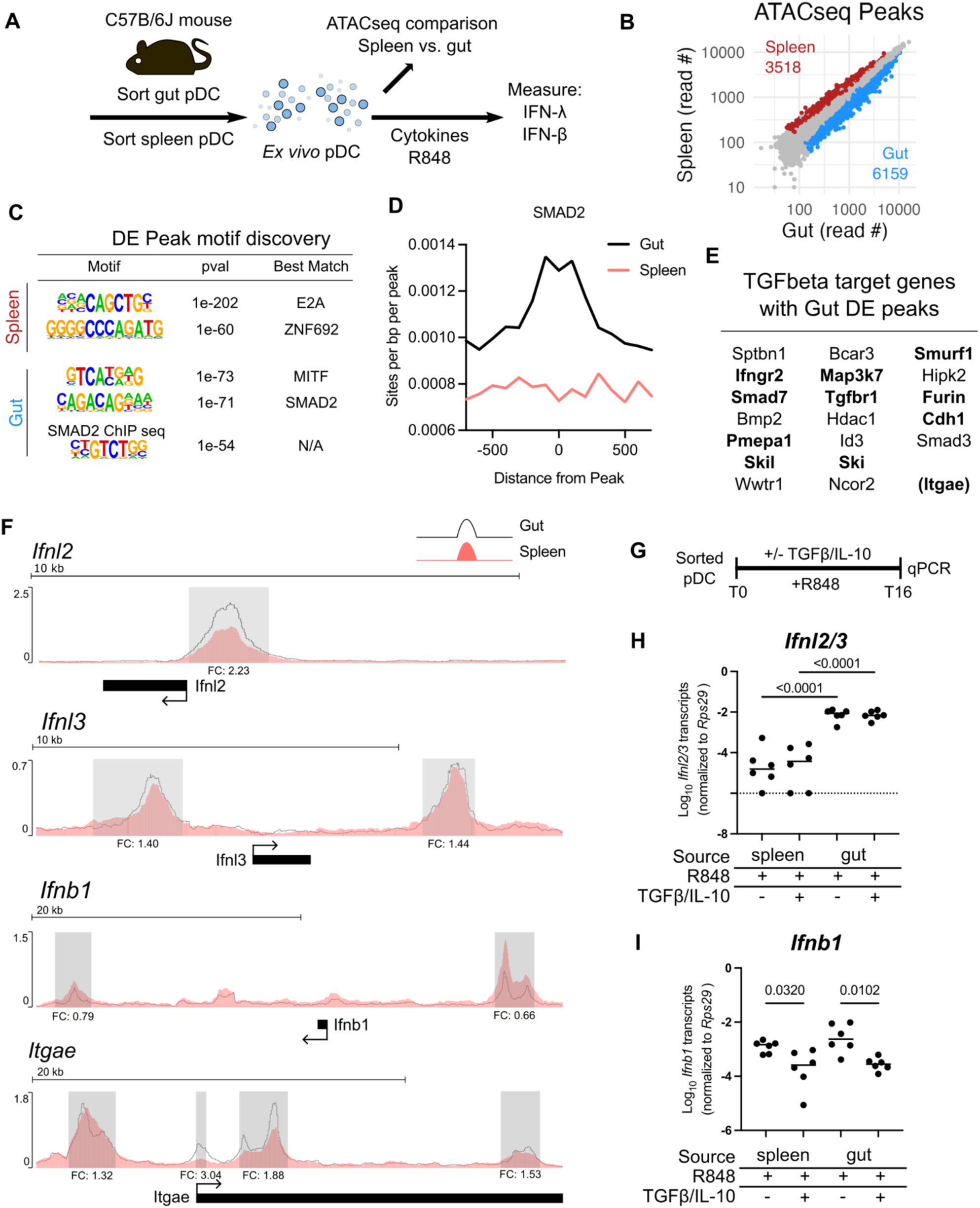
IFN type preference of intestinal pDC is associated with differential chromatin accessibility at SMAD2 binding sites. **A)**. Diagram of experiments on sorted mouse pDC from gut and splenic sources. **B).** Scatter plot showing differentially accessible regions for gut and spleen with padj < 0.05. **C).** De-novo motifs identified within gut-enriched and spleen-enriched peaks by HOMER and enrichment of a known SMAD2 motif. **D).** Plot showing the distribution and frequency of known SMAD2 binding motif within differentially accessible gut and spleen peaks. **E).** TGF-β HALLMARK pathway genes (*Itgae* is also included due to known TGF-β regulation) near peaks with increased accessibility in gut pDC. Genes with increased expression in scRNAseq data are in **bold**. **F).** Plots showing differential accessibility of gut (black line) and spleen (shaded salmon histogram) chromatin. Overlaid consensus ATAC-seq tracks for the indicated genomic loci from gut and spleen with open reading frames noted below and identified peaks indicated by shaded gray boxes. Fold-change of gut peaks relative to spleen peaks is listed under each peak region (FC). N=4 replicates for each tissue; genomic scale bars above tracks. **G).** Diagram of experimental design for stimulation of pDC *ex vivo*. **H-I)**. Expression of *Ifnl2/3* (**H**) and *Ifnb1* (**I**) from pDC isolated from gut or spleen of *n* = 6 mice from two experimental replicates and treated with the indicated cytokines and stimuli. Statistical significance by one-way ANOVA with Šídák’s multiple comparisons test.

In addition to the patterns of pDC regulation in the gut, we found that gut pDC had significantly increased accessibility at peaks near the transcription start sites (TSS) for *Ifnl2*, *Ifnl3*, and *Itgae* genes (**Fig. 5F**), consistent with their increased expression in our integrated scRNA-seq analysis (**Figs. 4E, 4G**). In contrast to IFN-λ genes, peaks ∼10kb upstream and downstream of the *Ifnb1* locus were significantly reduced in gut pDC relative to splenic pDC (**Fig. 5F**), consistent with reduced expression of this type I IFN gene in the gut. No peaks near other type I IFN genes (e.g. *Ifna4*) were significantly different between gut and splenic pDC. Together, the significant changes in genomic accessibility at IFN gene loci support a genomic basis for preferential IFN-λ production by pDC in the gut.

### TGF-β potentiates IFN-λ production

TGF-β, as well as interleukin 10 (IL-10), have been shown to suppress Type I IFN production by splenic pDC (Contractor et al., 2007). To test the relationship between genomic accessibility and cytokine stimulation in differential production of IFN types, we sorted pDC from gut and spleen as for ATAC-seq and cultured them *ex vivo* in the presence of the TLR7/8 agonist resiquimod (R848) with or without additional cytokine co-treatment (**Figs. 5A, 5G**). To minimize the time in culture *ex vivo*, we added R848 immediately after sorting, with or without TGF-β/IL-10 co-treatment, and collected cells after 16 hours for qPCR analysis. Under these conditions, we observed robust *Ifnl2/3* production by gut pDC with modest to undetectable levels in spleen pDC, regardless of TGF-β and IL-10 co-treatment (**Fig. 5H**). However, we observed that TGF-β/IL-10 co-treatment resulted in decreased *Ifnb1* stimulation in both spleen and gut pDC (**Fig. 5I**). These results support the previously reported role for TGF-β/IL-10 in repressing *Ifnb1* expression. Additionally, these data indicate that gut pDC intrinsically have a much greater capacity for *Ifnl2/3* production than spleen pDC which is unaffected by TGF-β/IL-10 co-treatment (**Fig. 5H**). The greater production of *Ifnl2/3* by R848-stimulated gut pDC is concordant with increased accessibility near TSS for these IFN-λ genes (**Fig. 5F**). Together these data indicate that increased accessibility at IFN-λ gene loci corresponds with substantially amplified production of *Ifnl2/3* upon stimulation.

Mouse pDC could be sorted in relatively low numbers from gut epithelium, limiting the experimental conditions in our IFN stimulation experiments (**Fig. 5G-I**). To perform a more detailed analysis of the effects of TGF-β and IL-10 and translate these findings to human cells, we used CAL-1 cells, a human pDC cell line established from a lymphoma patient (Maeda et al., 2005). This cell line is known to mount robust type I IFN response upon TLR7 stimulation with R848 (Joslyn et al., 2018). We pre-treated CAL-1 cells with TGF-β/IL-10, TGF-β alone, IL-10 alone, or media only for 16 hours and then stimulated IFN responses via R848 (**Fig. 6A**).

**Figure 6.**
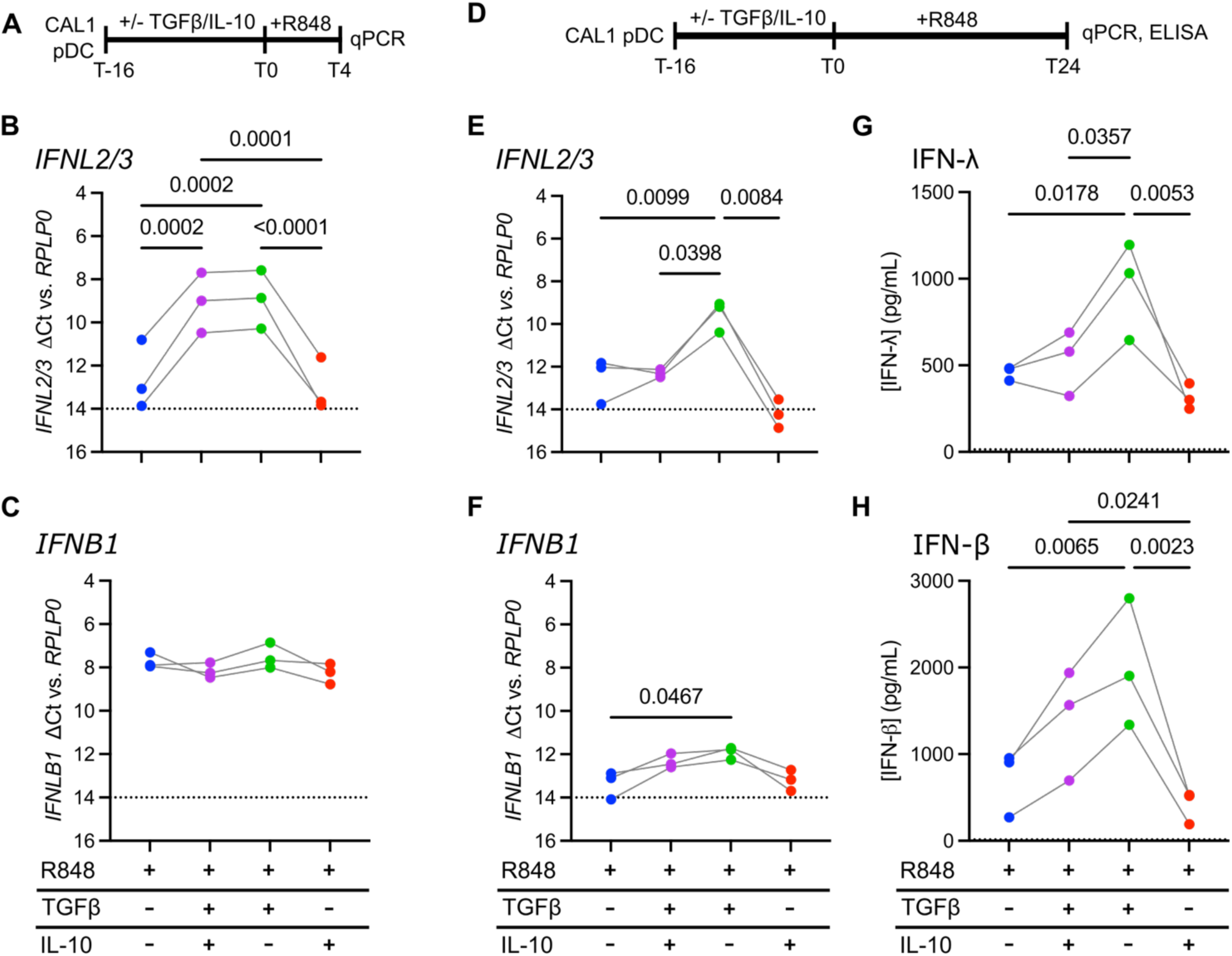
TGF-β potentiates IFN-λ production in CAL-1 human pDC. **A)**. Experimental design for 4-hour CAL-1 stimulation experiments. **B-C).** CAL-1 cells were treated with 10 ng/mL TGF-β and/or IL-10 for 16 hours in reduced-serum media, then stimulated with 1 µg/mL R848 for 4 hours prior to quantitating relative cycle threshold (delta Ct) for *IFNL2/3* (**B**) and *IFNB1* (**C**) relative to *RPLP0* by qPCR. **D).** Experimental design for 24-hour CAL-1 stimulation experiments. **E-H)**. CAL-1 cells were treated as in (A-C) but were stimulated for 24 hours. Relative cycle threshold number was determined by qPCR for *IFNL2/3* (**E**) and *IFNB1* (**F**), and IFN concentration in cell supernatants was determined by ELISA for IFN-λ (**G**) or IFN-β (**H**). All experiments were performed 3 times, and each data point is the mean of *n* = 3 technical replicates. Statistical significance determined by one-way repeated measures ANOVA with Tukey’s multiple comparisons test. Dashed lines for qPCR results (**B-C**, **E-F**) represent the highest detected quantity of the tested gene for unstimulated control cells. Dashed lines for ELISA results (**G-**H) represent the limit of detection based on the standard curve for the assay.

After 4 hours of R848 stimulation and no cytokine pre-treatment, there was minimally detectable *IFNL2/3*, but TGF-β or TGF-β/IL-10 pre-treatment resulted in a striking potentiation of *IFNL2/3* stimulation (**Fig. 6B**). In contrast, R848 stimulated substantial production of *IFNB1* at 4 hours with no statistically significant differences between any pre-treatment groups (**Fig. 6C**).

To assess longevity of IFN gene transcription and quantify secreted protein, we stimulated CAL-1 cells with R848 for 24 hours prior to collecting cell supernatants and RNA (**Fig. 6D**). We again observed a striking potentiation of *IFNL2/3* expression by TGF-β at this timepoint (**Fig. 6E**). However, TGF-β/IL-10 pre-treatment resulted in significantly less *IFNL2/3* expression compared to TGF-β alone at this timepoint (**Fig. 6E**), suggesting a dampening effect of IL-10 at longer times post-stimulation. In comparison, *IFNB1* was modestly expressed 24 hours after R848 stimulation in all groups (**Fig. 6F**), reflecting a rapid shutoff of this gene locus regardless of cytokine pre-treatment. TGF-β pre-treatment may delay this shutoff because it resulted in a small but significant increase in *IFNB1* at 24 hours (**Fig. 6F**). The potentiation of IFN gene transcription by TGF-β correlated with greater production of IFN proteins in supernatants at 24 hours post-stimulation (**Fig. 6G-H**).

These data from human CAL-1 cells support a model wherein TGF-β potentiates activation of the IFN-λ gene locus. Together with effects of IL-10 and other unknown signals within the intestinal milieu, TGF-β reprograms gut pDC to preferentially produce IFN-λ as befits the IFN response profile of the intestinal epithelium.

## Discussion

IFN-λ confers protection from enteric pathogens with reduced damaging effects on the host relative to Type I or Type II IFN (Wright and Nice, 2024; Stanifer et al., 2020). In the intestine, we observe that IFN-λ-dependent ISGs are expressed at homeostasis in a highly localized pattern, on the level of individual villi and small clusters of villi (**Fig. 1A-C**). Previous studies from our lab implicated an immune cell type in the production of IFN-λ in the mouse intestine through detection of *Ifnl2/3* transcripts in CD45^+^ cells from the epithelial fraction (Van Winkle et al., 2022). We also observed a reduced ISG response upon conditional deletion of the TLR adaptor gene myeloid differentiation primary response 88 (*Myd88*) in hematopoietic cells (Constant et al., 2025). In the current study, we established that a functionally, transcriptionally, and epigenetically distinct subset of pDC in the intestine are a primary contributor to IFN-λ production at homeostasis.

We previously found that localized ISGs correlate with local microbial abundance in the lumen (Constant et al., 2025). We hypothesize that a local concentration of luminal microbial products, together with infrequent transepithelial exposure to due to fluctuations in barrier strength, account for the localized pattern of ISG expression. The specific mechanism of transepithelial microbial exposure likely includes known routes of sampling (Knoop et al., 2013) and stimulation of TLR signaling by hematopoietic cells (Constant et al., 2025).

Here, we confirmed prior reports of pDC recruitment to the intestinal epithelium by microbial colonization and showed that they were more likely to be positioned within villi when microbial stimuli were present (**Fig. 2**). Thus, in addition to being a source of TLR ligands, microbial colonization drives signals that promote pDC recruitment towards the epithelium.

We showed that, among immune cells present in the intestine, only pDC make appreciable levels of IFN-λ at homeostasis. However, it is notable that the observed IFN levels are near the limit of detection (i.e., few IFN reads in **figure 4E**). Therefore, it is possible that we are missing contributory cellular sources of IFN that are less prolific than pDC. Indeed, the depletion of IFN-λ-producing pDC resulted in only a 50% reduction in homeostatic ISG expression in the epithelium (**Fig. 3B, 3F**). The residual epithelial ISG expression is likely resulting from other sources of IFN-λ, since this ISG expression is nearly absent in the IFN-λ cytokine knockout mice (**Fig. 1A-B**). T cells are reported to be capable of IFN-λ production (Swamy et al., 2015), and IECs preferentially produce IFN-λ upon stimulation or active viral infection (Odendall et al., 2014; Mahlakoiv et al., 2015; Van Winkle et al., 2022; Ingle et al., 2023). Our leading hypothesis for additional non-pDC sources of homeostatic IFN-λ is an IEC type; however, we suspect that low transcript abundance and a low proportion of IECs actively producing IFN-λ at homeostasis makes this technically challenging to verify without a model for conditional deletion of the entire IFN-λ gene locus.

We performed reciprocal bone marrow chimera experiments to determine if transplanted hematopoietic cells could restore homeostatic ISG signals in IFN-λ deficient mice (**Supplementary** Fig. 2). However, this experiment was not suitable for this purpose due to an elevated ISG signal in the intestinal epithelium of resulting chimeric mice (**Supplementary** Fig. 2B-C). IFN-λ plays a crucial role in intestinal healing after irradiation in allogeneic transplants (Henden et al., 2021) and can promote damage after irradiation in other contexts (Jena et al., 2024). It is likely that radio-resistant cells such as IECs produce elevated IFN-λ for at least six weeks in the absence of pDC-derived IFN-λ (*Ifnl2/3^-/-^* into WT CD45.1 group). However, given the incomplete chimerism in the CD3e^+^ compartment of bone marrow recipients (**Supplementary** Fig. 2G), intra-epithelial T cells are also a possible source of elevated IFN-λ after bone marrow transplant. These bone marrow chimera data emphasize that the specific cellular sources of IFN production in a healthy intestine are highly dependent on the context (e.g. sub-pathological inflammation). Nonetheless, our data clearly indicate that pDC play a key role in tonic IFN-λ signaling.

The finding that gut pDC preferentially produce IFN-λ over Type I IFN (**Fig. 1I, 4E**) raises the question of an underlying mechanism by which they are instructed. Prior data (Contractor et al., 2007) indicated that gut tissue cues such as TGF-β, IL-10, and PGE2 downregulate production of IFN-β by pDC, and we also observed suppression of *Ifnb1* transcript in sorted mouse pDC co-treated with TGF-β/IL-10/R848 relative to R848 stimulation alone (**Fig. 5I**). Increased CD103 expression on gut pDC relative to splenic pDC (**Fig. 1H, 4G**) may also contribute to suppression of Type I IFN in the gut. CD103 has been linked to restriction of Type I IFN expression in BM-derived dendritic cells (BMDCs) and cDC1 from *Itgae^-/-^* mice are more susceptible to viral infection (Duhan et al., 2021). However, suppressive factors that act on Type I IFN did not explain the increased production of IFN-λ by gut pDC (**Fig. 5H**).

Evidence of an active response to TGF-β in gut pDC was apparent from transcriptional and ATACseq analyses (**Fig. 4F-G, 5C-E**). Specifically, gut pDC had increased genomic accessibility relative to splenic pDC at peaks with SMAD2 binding motifs (**Fig. 5D**) including the TSS regions of IFN-λ genes (**Fig. 5F**). The correlation with gut pDC producing ∼16-fold more *Ifnl2/3* than splenic pDC (**Fig. 5H**) suggests that these epigenetic differences underlie enhanced IFN-λ production. Based on these data, our hypothesis is that TGF-β signaling drives increased capacity for IFN-λ production in gut pDC, while IL-10 together with other signals in gut (e.g. CD103 signaling) has inhibitory effects on IFN-β. We support this hypothesis and extend our findings to human cells through experiments with the pDC cell line CAL-1, wherein TGF-β pre-treatment substantially (∼8-fold) potentiates IFN-λ transcription (**Fig. 6**). In CAL-1 cells, Type I IFN is suppressed by IL-10 pre-treatment, but not TGF-β. It is possible that the timing of TGF-β treatment determines the effect on IFN expression. Our results with IL-10, however, are consistent with this cytokine’s canonical anti-inflammatory role and its specific previously known role as an inhibitor of Type I IFNs (Bruchhage et al., 2018; Duramad et al., 2003; Gary-Gouy et al., 2002).

In summary, we have identified that gut-intrinsic signaling reprograms pDC to preferentially produce IFN-λ over type I IFNs in mice and human cells. This altered IFN program is contrary to known pDC functions but is logical to allow pDC to signal effectively to IEC in the intestine without triggering widespread inflammation. This tissue-specific function of pDC as secretory sources of IFN-λ underlies our observations of highly localized ISG expression. We hypothesize that pDC are stimulated to produce IFN-λ after detection of microbial-associated molecular patterns originating in the lumen of the intestine, which have penetrated into the gut tissue by active or passive processes yet to be defined. The reprogramming of pDC which allows this compartmentalized signaling is largely dependent on TGF-β. These findings continue to reveal the plasticity of pDC and highlights their central role in innate immunity via production of IFNs.

## Materials and Methods

### Mice

Mice were housed in a specific pathogen-free facility at Oregon Health & Science university (OHSU). Procedures were approved by the Institutional Animal Care and Use Committee under protocol #IP00000228 in accordance with the standards provided in the Animal Welfare Act. All mice used in this study were maintained on the C57BL/6 background. Experimental groups were composed of littermate mice between 7 and 14 weeks of age and sexes were equally distributed among experimental groups. C57BL/6J WT and *Ifnl2/3*^-/-^ (*Ifnl3*^em1Mtba^) mouse strains were bred at OHSU and offspring used for experiments. CD45.1 congenic WT mice were purchased from Jackson Laboratories (stock #002014) and acclimated for at least one week at OHSU prior to use in experiments. BDCA2-DTR mice (Tg(CLEC4C-HBEGF)956Cln) were maintained at Washington University in St. Louis by the lab of Dr. Marco Colonna and bones were shipped to OHSU for use in bone marrow transplants.

### Bone marrow transplants

Bone marrow recipients were treated with split-dose 12 Gray radiation in an X-ray irradiator (RS2000, Rad Source Technologies). Two doses of 6 Gray were given approximately 4 hours apart. Mice were allowed to rest for 24 hours and administered 2.5×10^6^ live bone marrow-derived donor cells by retro-orbital injection. Mice were allowed to recover for 6-8 weeks prior to further experimentation.

### *In vivo* depletion of pDC

Mice were experimentally treated to reduce intestinal pDC as indicated in figure legends. Antibiotics were administered *ad libitum* for two weeks prior to sacrifice in autoclaved water (1 g/L ampicillin, 1 g/L metronidazole, 1 g/L neomycin, and 0.5 g/L vancomycin; Sigma, St Louis, MO). Anti-BST2 (clone 927) and isotype control (rat IgG2a, clone LTF2) antibodies (#BE0311 and #BE0090, Bio X cell, Lebanon, NH) were administered (2.5 mg/mL in 100 µL PBS) via retro-orbital injection at four and two days prior to sacrifice. Diptheria toxin (#150, List Labs, Campbell, CA) or sham (PBS) was administered via intraperitoneal injection at 20 ng/g over three consecutive days prior to sacrifice.

### RNA *in situ* hybridization

Intestinal tissue Swiss rolls were fixed in 10% neutral-buffered formalin (NBF) for 18–24 hr and embedded in paraffin. Tissue sections (5 μm) were cut and stored at room temperature with desiccant until staining. RNA *in situ* hybridization was performed using the RNAscope Multiplex Fluorescent v2 kit (Advanced Cell Diagnostics, Newark, CA) per protocol. As indicated in figure legends, slides were stained with anti-sense probes for *Ifit1* (ACD, #500071), *Epcam* (ACD, #418151), and *Siglech* (ACD, #432671). Slides were counterstained with DAPI, mounted with ProLong Gold antifade reagent (ThermoFisher), and imaged using a Zeiss ApoTome2 on an Axio Imager, with a Zeiss AxioCam 506 (Zeiss). Collected images were batch processed in Zeiss Zen 3.1 using unstained control slides to set background values. Fluorescent signal was quantified using Zeiss Zen 3.1 as positive fluorescent area for each target relative to the total fluorescent area of the tissue section as determined by DAPI. *Siglech* signal was used to identify pDC.

### Epithelial strip and CD45 enrichment

Epithelial fractions were prepared by non-enzymatic dissociation as previously described (Nice and Constant, 2024; Van Winkle et al., 2022). Briefly, mouse ileum was longitudinally opened and agitated by shaking in stripping buffer (5% fetal bovine serum (FBS), 5 mM EDTA, 1 mM dithiothreitol, 1X penicillin-streptomycin-L-glutamine, in PBS) for 20 min at 37°C. Epithelial stripping solution was filtered through a 100 μm cell strainer and dissociated cells were collected for use in qPCR analysis, flow cytometry, and magnetic bead enrichment.

### Quantitative PCR

RNA from stripped IECs was isolated with TRIzol (Life Technologies, Carlsbad, CA) per the manufacturer’s protocol. RNA was isolated from magnet-enriched and FACS-sorted cells with a Zymo Quick-RNA Viral Kit (Zymo Research, Irvine, CA). RNA (larger of 1 μg or 5 μL) was treated with the DNAfree kit (Life Technologies) and used as a template for cDNA synthesis (ImProm-II reverse transcriptase system, Promega, Madison, WI).

Quantitative PCR for mouse target genes was performed (PerfeCTa qPCR FastMix II, QuantaBio, Beverly, MA) and absolute transcript copy numbers were determined using standard curves generated from synthetic gBlocks (IDT) containing target sequences. Absolute copy numbers were normalized to gene copies of ribosomal protein S29 (*Rps29*). The following primer-probe assays for selected genes were ordered from IDT (Coralville, IA): *Rps29* (Mm.PT.58.21577577), *Ifit1* (Mm.PT.58.32674307). Primer-probe assays for *Ifnl2/3* and *Ifnb1* were previously designed (Van Winkle et al., 2020) with the following sequences: *Ifnl2/3* (Primer 1 – GTTCTCCCAGACCTTCAGG, Primer 2 – CCTGGGACCTGAAGCAG, Probe – CCTTGCAGGCTGAGGTGGC); *Ifnb1* (Primer 1 – CTCCAGCTCCAAGAAAGGAC, Primer 2 – GCCCTGTAGGTGAGGTTGAT, Probe – CAGGAGCTCCTGGAGCAGCTGA). For CAL-1 human pDC cell experiments, approximately 1×10^6^ cells were lysed in 0.75 mL TRIzol. RNA was isolated by the Direct-zol kit (Zymo Research, Irvine, CA) and 0.5 μg was prepared for qPCR as for other samples. Human target genes were quantified relative to the cycle threshold of the ribosomal protein lateral stalk subunit P0 (*RPLP0*) by the delta Ct method. Predesigned primer-probe assays were used for *RPLP0* (Hs.PT.39a.22214824) and *IFNB1* (Hs.PT.58.39481063.g), and an assay was designed for *IFNL2/3* with the following sequences: Primer 1 – GCGACTCTTCTAAGGCATCTT, Primer 2 – CCACATAGCCCAGTTCAAGT, Probe – ACAGGAGCTGCAGGCCTTTAAGAG.

### ELISA

ELISA assays were performed to quantify IFN-λ and IFN-β in CAL-1 cell supernatants. The DuoSet human IFN-β ELISA kit (R&D Systems, Minneapolis, MN; DY814) was used per manufacturer recommendations. The VeriKine-DIY human IFN-Lambda 3/1/2 (IL-28/29/28A) ELISA kit (PBL Assay Science, Piscataway, NJ; 61840) was used per manufacturer recommendations, with the following modifications: capture antibody was diluted to 2.0 µg/mL working concentration, detection antibody was diluted to 800 ng/mL, and signal was detected with high-sensitivity TMB substrate (BioLegend; 421501).

### Flow cytometry and magnetic bead enrichment

Dissociated IECs and splenocytes were collected and stained for flow cytometry. Cells were stained with the following reagents and antibodies, conjugated to fluorophores indicated in **Supplementary Table 1D** and, where appropriate, figures and associated legends: Zombie Aqua viability dye (BioLegend), DAPI, Fc receptor-blocking antibody (CD16/CD32; clone 93; BioLegend), anti-CD45 (clones 30-F11 or I3/2.3; BioLegend), anti-CD45.1 (clone A20; BioLegend), anti-CD45.2 (clone 104; BioLegend), anti-CD3e (clone 145-2C11; BioLegend), anti-CD11b (clone M1/70; BioLegend), anti-CD11c (clone N418; BioLegend), anti-CD103 (2E7; BioLegend), anti-CX_3_CR1 (clone SA011F11; BioLegend), anti-IA/IE (clone M5/114.15.2; BioLegend), anti-Ly6C (clone HK1.4; BioLegend). Where indicated, dissociated cells were enriched using MojoSort Mouse anti-APC Nanobeads (BioLegend, #480072) after flow cytometry staining for anti-CD45 of anti-CD45 and anti-SiglecH with APC fluorophores following manufacturer protocols. All data were analyzed using FlowJo software (BD Biosciences). Gates were set based on unstained and single-fluorophore stains.

### *Ex vivo* culture of mouse pDC

Splenocytes were obtained by manually dissociating spleen tissue through a 70 μm cell strainer followed by red blood cell lysis with ACK buffer. Stripped intestinal epithelial fraction was isolated from small intestine and CD45^+^ SiglecH^+^ double-positive cells were enriched as described above. Splenocytes and epithelium-associated leukocytes from individual mice were stained and sorted as indicated in **Supplementary** Figure 3 on a FACSAria Fusion (Beckton Dickinson) using a 70 μm nozzle directly into R10 media (RPMI media supplemented with 10% FBS, 1X non-essential amino acids, 1 mM sodium pyruvate, 10 mM HEPES, and 1X penicillin-streptomycin-L-glutamine). Approximately 5,000-15,000 cells per sample were sorted, concentrated, and cultured in duplicate wells (approx. 2,500-7,500 cells/well) of a non-tissue culture treated U-bottom 96 well plate in 50-60 μL R10 media with or without R848 (10 µg/mL; InvivoGen, San Diego, CA; tlrl-r848), IL-10 (100 ng/mL; ProteinTech, Rosemont, IL; HZ-1145), or TGF-β (100 ng/mL; ProteinTech, Rosemont, IL; HZ-1011) as indicated in figure legends. Cells were recovered from culture media and cDNA libraries were prepared using the SuperScript IV Low-Input cDNA synthesis kit (ThermoFisher) using the manufacturers protocol with the following modifications: reverse transcription was performed for 60 minutes, cDNA pre-amplification was performed for 12 cycles, and all libraries were purified prior to analysis by qPCR using Ampure XP beads (Beckton Dickson).

### CAL-1 cells

CAL-1 cells were grown in complete RPMI (Roswell Park Memorial Institute media, supplemented with 1X non-essential amino acids, 1 mM sodium pyruvate, 10 mM HEPES, and 1X penicillin-streptomycin-L-glutamine) with either 10% FBS (R10) for standard growth conditions or 0.1% FBS (R0.1) for serum starvation. For all experiments, approximately triplicate wells with 1.3×10^6^ cells were serum-starved for 16 hours prior to stimulation. As indicated in figure legends, cells were treated with 10 ng/mL TGF-β and/or IL-10 in R0.1, then stimulated with 1 µg/mL R848. Assay sensitivity for gene expression and ELISA measurements was determined using unstimulated control samples.

### Single-cell RNAseq

Epithelium-associated leukocytes were isolated as described above from four male and four female mice. Myeloid subsets were sorted on either an Influx or FACSAria Fusion (Beckton Dickinson) using the gating strategy shown in **Supplementary** Figure 1A. In addition to fluorescent antibodies, cells were stained with antibody-derived tags (ADT) and cells from each mouse were stained with a different hashtag oligonucleotide (HTO), listed in **Supplementary Table 1A**. Pooled cells from all mice were prepared using 10X Genomics Single Cell 3’ V3 reagents into gene expression (GEX), ADT, and HTO libraries and sequenced using the Illumina NovaSeq platform.

### Single-cell RNAseq analysis

Reads were mapped to the mouse transcriptome (mm10-2020-A) and cells were filtered using the cellranger v7.0.0 pipeline and resulting filtered feature barcode matrix was loaded into Seurat version 4.1.3 for analysis. Samples were de-multiplexed using the HTODemux function on HTO count data and singlets were retained for further analysis. Data was further subset to retain cells with between 200-4000 features and less than five percent mitochondrial reads. GEX data was then scaled, normalized, and clustered using 500 variable features, dims = 1:12, and resolution = 0.15. Clusters were defined based on GEX and ADT data comparison to known cell lineage markers. Cells within the pDC cluster from figure 1 were exported as a counts matrix for integration with public pDC datasets in figure 4.

Data integration in figure 4 was performed using Seurat version 5.0.3 in R version 4.3.3. The following counts matrices were used to generate Seurat objects: 1) gut pDC from our dataset GSE304885, 2) Bone marrow pDC from GSE114313, 3) spleen pDC from GSE114313, and 4) spleen pDC from GSE196720. Datasets were prefiltered according to the parameters outlined in **Supplementary Table 1B**. The data were then normalized, variable features were identified, and data were integrated using 5000 anchors and 50 dimensions. Integrated data were then clustered using following functions: ScaleData, RunPCA, FindNeighbors (dims = 1:8), FindClusters (resolution = .1), and RunUMAP (dims = 1:8). Marker genes for cell clusters were identified by joining the RNA layers (JoinLayers function) and using FindAllMarkers function with restriction of the feature list to genes detected in all four datasets. Additionally, all datasets were downsampled to 200 cells (size of smallest dataset) to avoid domination of the comparison by larger datasets. Marker genes presented in the original publications of these data were used to infer identity of integrated clusters and dotplots were generated with downsampling of datasets to 200 cells. A ranked gene list for geneset enrichment was generated using the FindMarkers function to compare gut cells to concatenated spleen 1 and spleen 2 cells within the pDC cluster. Genes were ranked (gut vs spleen) based on signed z-score, and enrichment of HALLMARK pathways (mSigDB.org) within the ranked list was determined using the fgsea package.

### ATAC-seq and analysis

Sorted pDC were isolated as for *ex vivo* culture (**Supplementary** Figure 3) and immediately processed for ATAC-seq according to manufacturer specifications using the Fixed Cell ATAC-Seq Kit (Active Motif, #53151).

Resultant libraries were sequenced on a NovaSeq 6000 (Illumina). Sequencing data was processed using the NextFlow NF-Core ATAC-Seq pipeline (Ewels et al., 2020; Patel et al.), version 25.04.6, build 5954 (01-07-2025 11:27 UTC). Quality assessment metrics are outlined in **Supplementary Table 1E**. Motif enrichment analysis was performed in HOMER version 5.1 (Heinz et al., 2010). ATAC-seq tracks were visualized and prepared for publication using the Integrative Genomics Viewer (Thorvaldsdóttir et al., 2013; Robinson et al., 2011).

## Data availability

Sequencing data generated for this project is available from the gene expression omnibus (GEO) under parent accession number GSE304886. Raw fastq files and processed data files for RNAseq and ATACseq are available from subsidiary accession numbers GSE304885 and GSE304884 respectively. All other data and reagents are available upon request from the lead contact.

## Acknowledgements

We would like to thank Dr. Patrick Flynn and Dr. Evan Lind for their generous gift of antibody-derived tags to identify immune cells in RNA seq, Dr. Marina Cella and Dr. Marco Colonna for procuring BDCA2-DTR mouse bone marrow, and Dr. Helene Jahn for providing insightful critique of the manuscript. Illumina sequencing was performed by the OHSU Massively Parallel Sequencing Shared Resource, which is funded in part by the NCI Cancer Center Support Grant P30CA069533 awarded to the OHSU Knight Cancer Institute. We also thank the following core facilities for providing technical support and instrumentation on this project: the OHSU Advanced Light Microscopy core (RRID: SCR_009961), the OHSU Histopathology Shared Resource, and the OHSU Flow Cytometry Core. Funding for this work was provided by National Institutes of Health grants AI130055 (TJN), T32-AI007472 (JVW), and T32-GM142619 (KAM).

## Author contributions

Conceptualization: DAC, JVW, TJN Methodology: DAC, JVW, TJN, PAN, GM

Investigation: DAC, JVW, TJN, PAN, GM, KAM, MEL, AG, SRM

Visualization: DAC, TJN, JVW, PAN Funding acquisition: TJN

Project administration: TJN, DAC Resources: RS, PFR

Data curation: DAC, TJN Formal analysis: DAC, TJN Supervision: TJN, DAC, RS Writing – original draft: DAC

Writing – review & editing: DAC, TJN, JVW, PAN, KAM, SRM, AG

## Supplementary materials

**Supplementary figure 1.** Representative flow cytometry gating strategies for sorted cells used in Figure 1.

**Supplementary figure 2.** Radiation-resistant cells can stimulate localized IFN-λ-dependent gene expression in the context of bone marrow transplantation

**Supplementary figure 3.** Representative gating of defined single cells used to sort pDC from mouse intestine

**Supplementary file 1.** Spreadsheet containing supplemental tables 1A-1E.

**Supplemental table 1A.** Antibodies used in preparation of scRNAseq libraries.

**Supplemental table 1B.** Prefilters used for data integrated and re-analyzed in figure 4.

**Supplemental table 1C.** HALLMARK pathways in gene set enrichment analysis of integrated data in figure 4.

**Supplemental table 1D.** Fluorescent antibodies used in flow cytometry experiments.

**Supplemental table 1E.** Quality assessment metrics for ATACseq data in figure 5.

