## Supplementary Figures for "Plasmacytoid dendritic cells in the intestine preferentially produce interferon lambda at homeostasis contributing to tonic localized innate immune responses"

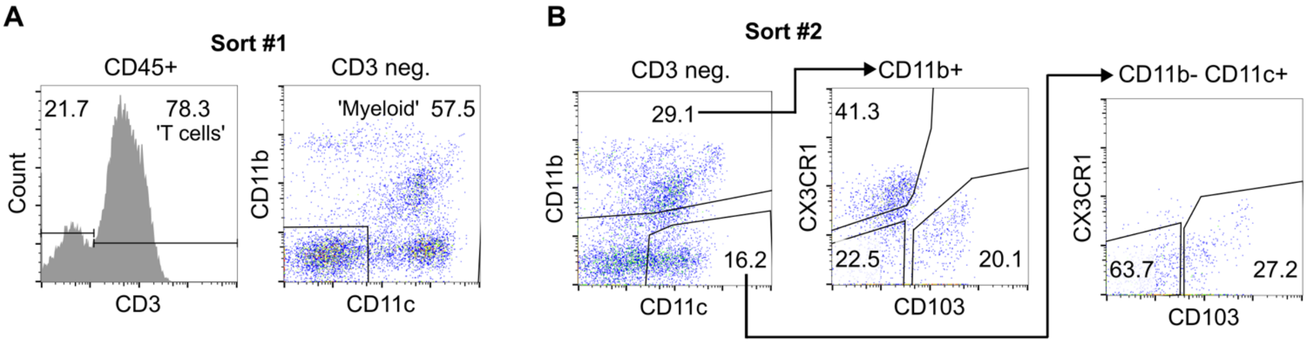


**Supplementary figure 1. Representative flow cytometry gating strategies for sorted cells used in Figure 1. A, B.** Representative FACS plots depicting gating for cells based on expression of CD3, CD11c, CD11b (A and B), $CX_3CR1$, and CD103 (B).

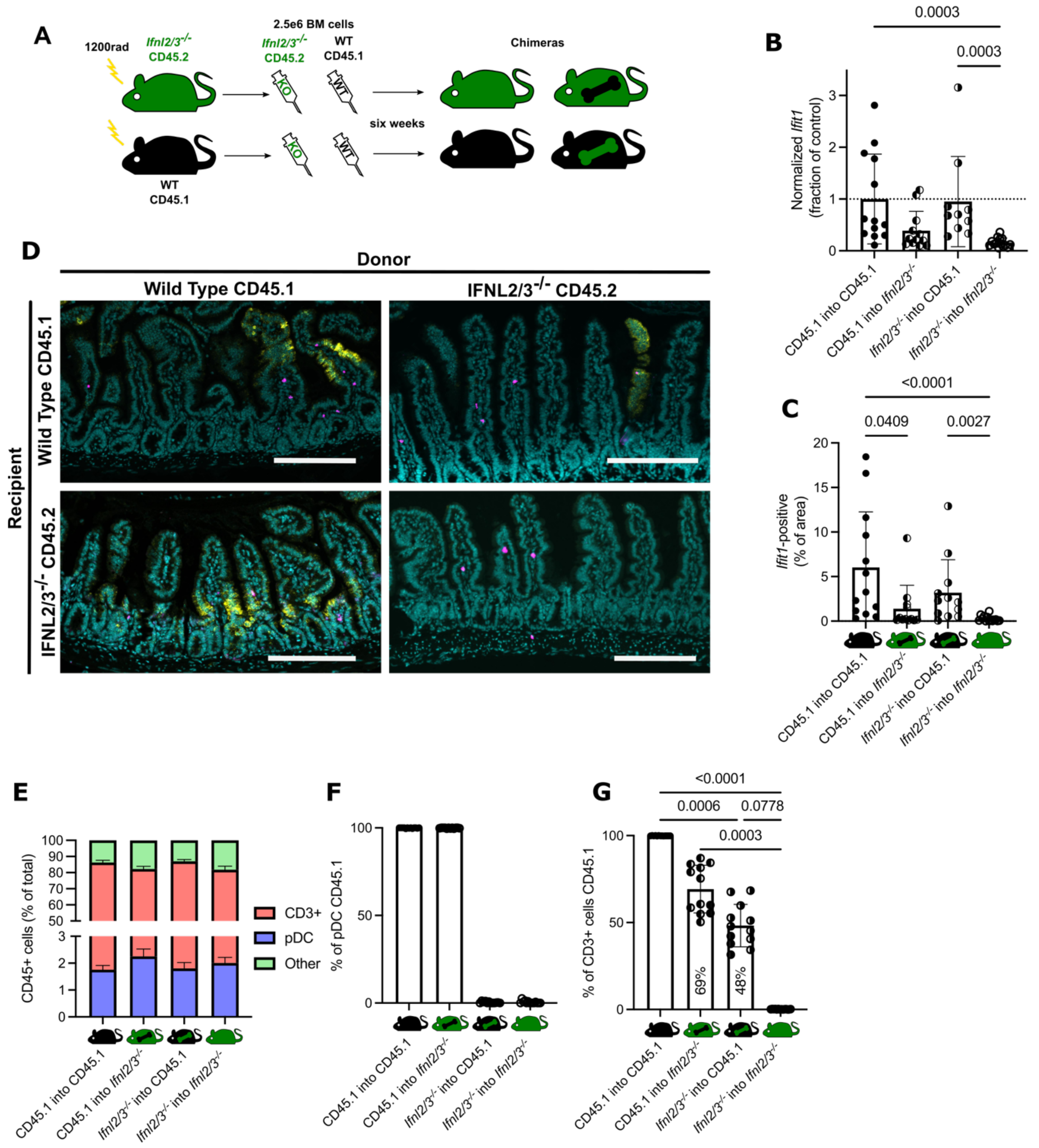


**Supplementary figure 2. Radiation-resistant cells can stimulate localized IFN-λ-dependent gene expression in the context of bone marrow transplantation. A.** Diagram of reciprocal bone marrow transfers of CD45.2 *Ifnl2/3*$^{-/-}$ and CD45.1 wild-type mice. **B.** Expression of *Ifit1* as measured by qPCR. **C.** Quantitation of intestinal area positive for *Ifit1* expression by RNAscope in indicated mouse strains. **D.** Representative images from RNAscope experiments. Scale bars = 50 µm. **E.** Percentage of cell types identified by flow cytometry in stripped intestinal epithelia of bone marrow chimeras. **F**. Percentage of pDC identified as CD45.1 (WT). **G.** Percentage of CD3$^{+}$ cells identified as CD45.1 (WT). Data for each reciprocal chimeric group was collected from three independent experiments with total *n* = 14 *Ifnl2/3*$^{-/-}$ into *Ifnl2/3*$^{-/-}$, *n* = 12 *Ifnl2/3*$^{-/-}$ into WT, *n* = 12 WT into *Ifnl2/3*$^{-/-}$, and *n* = 13 WT into WT. Statistical significance by Kruskal-Wallis test with Dunn's multiple comparison test in (B, C, G).

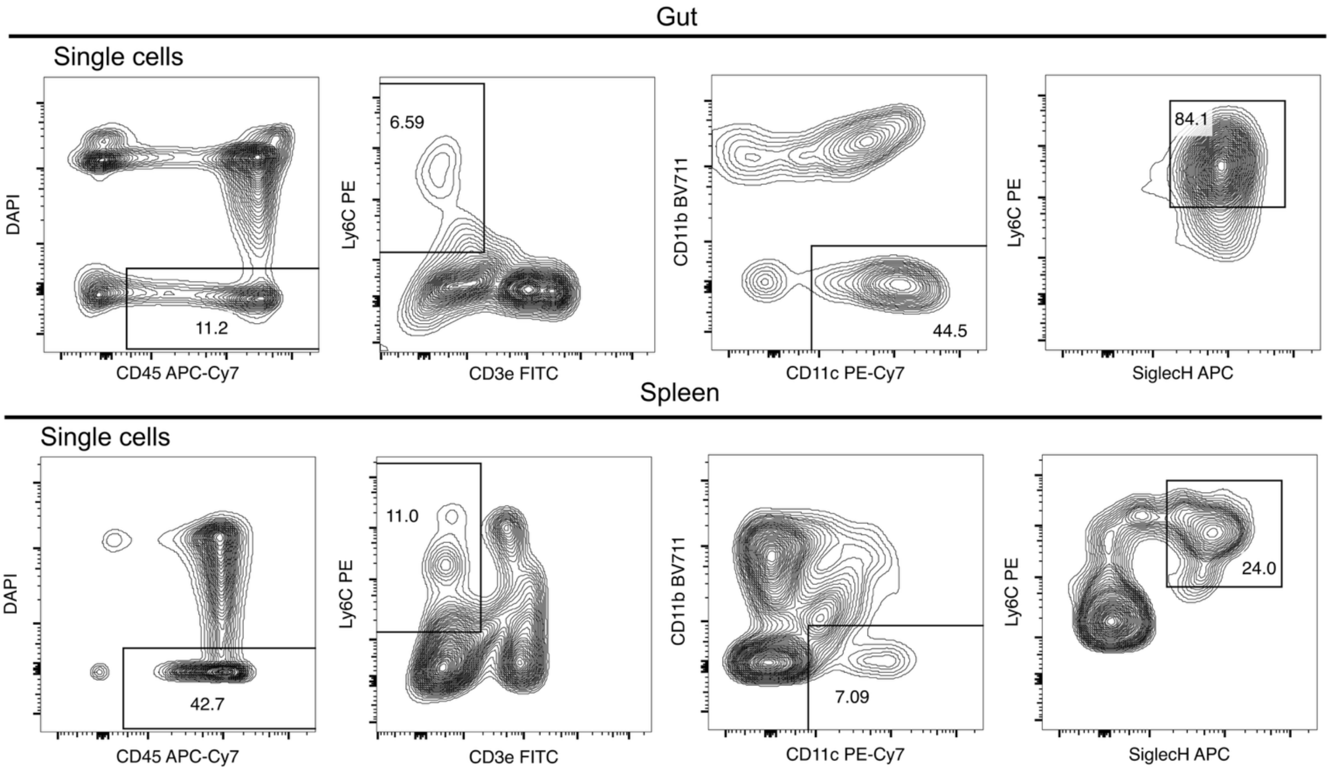


**Supplemental Figure 3.** Representative gating of defined single cells used to sort pDC from mouse intestine (Gut, top) and spleen (bottom) for experiments in Figure 5. Numbers in gates represent percentages of parent population.
